# Pan1c : a pipeline to easily build chromosome-level pangenome graphs

**DOI:** 10.64898/2026.04.17.719212

**Authors:** Alexis Mergez, Martin Racoupeau, Philippe Bardou, Benjamin Linard, Fabrice Legeai, Frédéric Choulet, Christine Gaspin, Christophe Klopp

## Abstract

The advances of sequencing technologies and the availability of high-quality genome assemblies for many genotypes per species, give the opportunity to improve sequence alignment rate and quality, and the variant calling accuracy by including all genomic variations in a graph reference, called a pangenome graph. Because the process of building and analysing a pangenome graph is still complex, with related software packages under development, there is an important need for releasing user-friendly pipelines for this emerging research area. `Pan1C` is a pipeline based on a chromosome-by-chromosome graph construction strategy. It integrates two complementary strategies for building pangenomes and produces informative metric plots and graphics using a large set of tools. By benchmarking `Pan1C` on human, fungal, and wheat assemblies, which span a wide range of genome sizes and complexities, we showed the interest of `Pan1C` for assembly and graph validation as well as for performing primary analyses.

## 1. Introduction

Advances in genome sequencing technologies have made it feasible for many laboratories to generate near-complete chromosomal assemblies of large genomes [3]. For species studied by large research communities, dozens of high-quality assemblies are now publicly available. In particular, species of agronomic interest are represented in international databases by extensive sets of breed- or cultivar-specific assemblies. A powerful way to exploit these datasets is the construction of pangenome graphs, which are increasingly replacing reference-based pairwise alignments. By capturing the genomic diversity present within a population, pangenome graphs can substantially improve analytical accuracy and completeness [25] [24] [17]. Two main tools have emerged for building such graphs, each based on a different methodology. The PanGenome Graph Builder (`PGGB`) relies on all-vs-all alignments [8] while `Minigraph-Cactus` (`MC`) pipeline uses reference-based alignments at chromosome-scale [13]. The two approaches produce topologically different graphs [2] [6] and differ on computation time with `PGGB` being significantly more time-consuming [2] and sometimes impossible to run. Aside from the graph-building tools, comprehensive pangenome toolkits such as `vg` [9] and `ODGI` [11] have emerged to support graph manipulation and downstream analyses. They include extensive sets of functionalities, specific file formats and options that are not easily accessible to non-expert users. To facilitate routine analyses, it is thus essential to provide documented, user-friendly workflows and to generate outputs that are easily interpretable, including graphical summaries that allow users to quickly and readily assess results.

Several workflows have emerged to facilitate pangenome graph construction, namely `nf-core/pangenome` [12] and `MoGAAAP` [22]. The first focuses on optimising pangenome graph construction, leveraging High Performance Computing (HPC). Sequence alignment of `PGGB` is distributed across multiple nodes, and every steps of the workflow is parallelised to improve overall computation time and efficiency. The workflow also allows to build chromosome-community graphs by clustering similar input sequences, simplifying the final graph into biological relevant components. Yet, this grouping does not ensure unique chromosome components and comparison between assemblies. MoGAAAP focuses on haplotype assembly before pangenome construction. Using raw input reads, the workflow outputs high-quality assembly draft, assembled into chromosomes using a high-quality reference assembly. Assembly standardisation eases gene-based pangenome graph construction and analysis. Both workflows produces quality assessment reports, yet in case of `nf-core/pangenome`, these can be limited as input quality is not checked and can directly impacts the quality of the pangenome.

Hereafter we present `Pan1C` a modular, scalable, and reproducible workflow for building and analysing pangenome graphs. `Pan1C` uses a chromosome-by-chromosome strategy which not only contributes to reduce overall computation time but also enhances interpretability. By constructing each chromosome graph independently, this strategy breaks the global complexity into manageable, biologically relevant units that can be more easily validated thanks to meaningful metrics and graphical plots generated. Quality and observed variations of both input assemblies and pangenome graphs are assessed and compared to improve downstream analysis.

## 2. Materials and Methods

### 2.1. Implementation and configuration

`Pan1C` is implemented in `Snakemake` [14], and uses standard YAML configuration files to define data paths and tool-specific options, allowing fine-grained control of graph construction. `Snakemake` also handles job scheduling and memory allocation while the workflow configuration specifies the maximum amount of preemptible resources. Reproducibility and ease of deployment are ensured through `Conda` and four version-controlled, upgradable, `apptainer` [15] images bundling softwares and dependencies. `Pan1C` was developed for `Slurm`-based highperformance computing (HPC) systems but has also been tested on standalone machines and non-`Slurm` HPC environments. The workflow execution can be monitored through independent rule log files as well as a global log file summarising the run. The `Pan1C` GitLab group (https://forge.inrae.fr/genotoul-bioinfo/Pan1C) contains source codes, container recipes, data download information, a complete documentation, a tutorial and usage examples. The `Pan1C-View` interface is implemented in HTML, CSS, and JavaScript, and uses a set of JSON files generated by the workflow to dynamically load and display tables and graphical outputs. The interface can be opened locally (by launching a lightweight HTTP server) or hosted on any standard web server, the code being provided as fully self-contained within a single .tar.gz archive (see our gitlab repository for more information).

### 2.2. Datasets

Three datasets corresponding to species with different sizes and complexities were chosen in order to illustrate representative cases in which `Pan1C` can be used. Human genomic data were chosen because they combine high assembly quality and well-characterised genomic variations and have been used by Andreace and colleagues in [2] to benchmark pangenome graph representations. We used 2 haplotypes from 4 different individuals of dataset 2 and the CHM13 and GRCh38 (HG00438) synthetic references. We discarded chromosome Y of CHM13 *Homo sapiens* reference assembly since it was not present in all assemblies. Constructing a pangenome graph for bread wheat was also considered as valuable for a test case due to the species genomic complexity and diversity. Indeed, bread wheat is an allohexaploid with a very large genome (∼15 Gb) that contains extensive structural variations and an extreme level of repetitive sequences [19], transposable elements representing 85% of the sequence [23]. Assemblies of 4 *Triticum aestivum* (bread wheat) cultivars including the reference IWGSC_RefSeq_v2.1 (GCA_018294505.1) were downloaded from URGI servers and NCBI respectively. *Aspergillus fumigatus* is a major human fungal pathogen, and its adaptability is driven by variations in genes and loci associated with virulence, antifungal resistance, environmental survival, and metabolic flexibility. One hundred *Aspergillus fumigatus* haplotypes were downloaded from NCBI and used to test `Pan1C` ability to handle large numbers of small-size genomes.

The main features of the 3 datasets are listed in Table 1. More complete information is given in supplementay tables 1-3 including accession numbers and N50.

**Table 1.** Main characteristics of datasets.

| Species | Reference | #individuals | #assemblies | Mean size (Mb) |
| --- | --- | --- | --- | --- |
| *Aspergillus fumigatus* | GCA_000002655 | 100 | 100 | 29 |
| *Homo sapiens* | GCA_009914755 (CHM13) | 6 | 10 | 3,005 |
| *Triticum aestivum* | GCA_018294505.1 | 4 | 4 | 14,208 |

### 2.3. Preparing input

`Pan1C` requires a set of compressed genome assemblies (.fa.gz) as input, including a highquality chromosome-level reference assembly. Depending on the quality of the input assemblies, different preprocessing steps are required (see cases and preprocessing steps at https://forge.inrae.fr/genotoul-bioinfo/Pan1c/pan1c/-/wikis/home/Tutorial). The reference assembly is used to scaffold non-chromosome-level assemblies into chromosomes using `RagTag` [1]. Briefly, `RagTag` only orders contigs against a reference, but does not modify the sequence itself. Chromosome-level input assemblies avoid this step, but require a table to map chromosome IDs to their reference equivalent. Optional sanity-check scripts are provided to check all input files, prepare the configuration file and generate the script for both local and `Slurm` cluster execution.

## 3. Results

### 3.1. The Pan1C workflow

The `Pan1C` workflow metro-map is shown in Fig. 1. It is divided into three main steps. During pre-processing (Fig. 1-A), a script prepares and checks inputs, chromosome sequences are clustered, assembly metrics are computed using `QUAST` [18] and `Assemblathon-stats` [4]. `SyRI` [10] is used to detect synteny and variations between haplotypes that are expected in the final graph. Quality control is essential as the overall quality of a pangenome strongly depends on the quality of its haplotype-resolved chromosome-level genome assemblies. In the main step (Fig. 1-B), chromosome-graphs are built in parallel using `PGGB` [13] and/or `MC` [8]. In the final step (Fig. 1-C), per-chromosome graphs are merged into per-tool graphs using `odgi merge` [11]. `Panacus` [20], as well as a custom script are used to generate various graph statistics, including node, edge and base pair counts, shared length, etc… `Pan1C` enables the visualisation of distances between chromosomes. To compute these distances, we first generate, for each chromosome, a table reporting the proportion of sequence shared between all pairs of paths. This produces the “Shared content” tables using the “Shared.prop” metric. For each pair of chromosomes, we then compute the Frobenius norm between their corresponding tables, yielding a similarity score that reflects differences in the amount of shared regions between chromosomes. Consequently, a large distance between two chromosomes indicates substantial differences between their sharing matrices, which may point to an anomaly, such as a potential assembly error.

**Figure 1.**
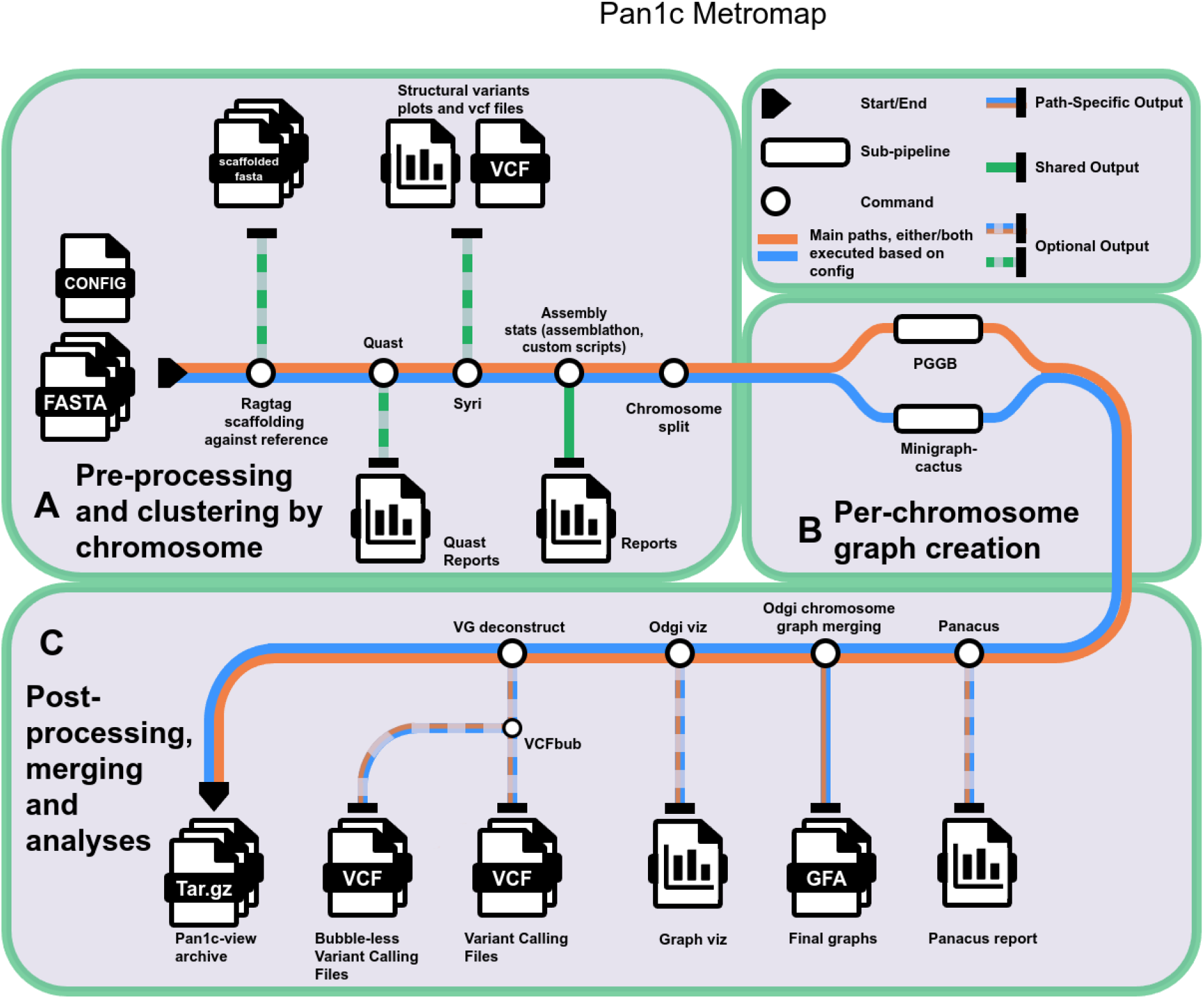
`Pan1C` workflow metromap. Colored paths represent different data flows between tools and outputs. Paths that diverge from the main route (green) represent optional steps. (A) Pre-processing and clustering by chromosomes. (B) Graph construction. (C) Per-chromosome graph merging and generation of graph-based outputs.

Optionally, `VG deconstruct` [16] can be used to call variants using the reference assembly, producing a standard Variant Call Format (VCF) file. For visual analysis, `odgi viz` is called to produce a linear representations showing synteny and structural variants between haplotypes. All resulting data files useful for downstream analyses are gathered in a single directory. Finally, the pipeline generates an archive ready for an interactive exploration of the pangenome graph in `Pan1C-View`.

### 3.2. Pan1C-View

Pan1C-View is an interactive interface designed to summarise and visualise the results produced by `Pan1C`. It allows users to explore assembly quality, pangenome graph consistency, and structural variation patterns without requiring any additional software installation. It is organised into four main sections. The **Summary** section provides dataset information, software versions and configuration details to insure reproducibility. It also includes download links to the generated .GFA (version 1.0), .XG (`VG` indexed file format) and .VCF files. GFA files include in their header all parameters used to generate them. The **Assemblies** section displays a table of Assemblathon [5] metrics, allowing users to verify that all assemblies meet expected quality. Heatmaps of chromosome lengths Fig. 2-B and visualisations such as contig position plots Fig. 2-A and `SyRI` plots (per chromosome and per haplotype) are available to visually assess structural similarities and differences between assemblies. `SyRI` plots can also be used to assess the coherence of syntenic regions and structural variants represented in the graph. The **Graphs** section presents a table of the general metrics of the graph and multiple plots for both `PGGB` and `MC`, enabling comparison of the two graph-construction methods. The path composition barplot shows how the graph is composed in terms of shared and assembly-specific nodes. Heatmaps show chromosome-to-chromosome distance (Fig. 2-C) and shared sequence content (Fig. 2-D) while 2D `ODGI` visualisations provide intuitive exploration of graph topology and structural variation. Finally, a `Panacus` report is also included, summarising pangenome growth trends. The **Variants** section includes a barplot showing the number of insertion/deletion (INS/DEL) per assembly and a plot depicting the INS/DEL size distribution extracted using `VG deconstruct` and `SyRI`.

**Figure 2.**
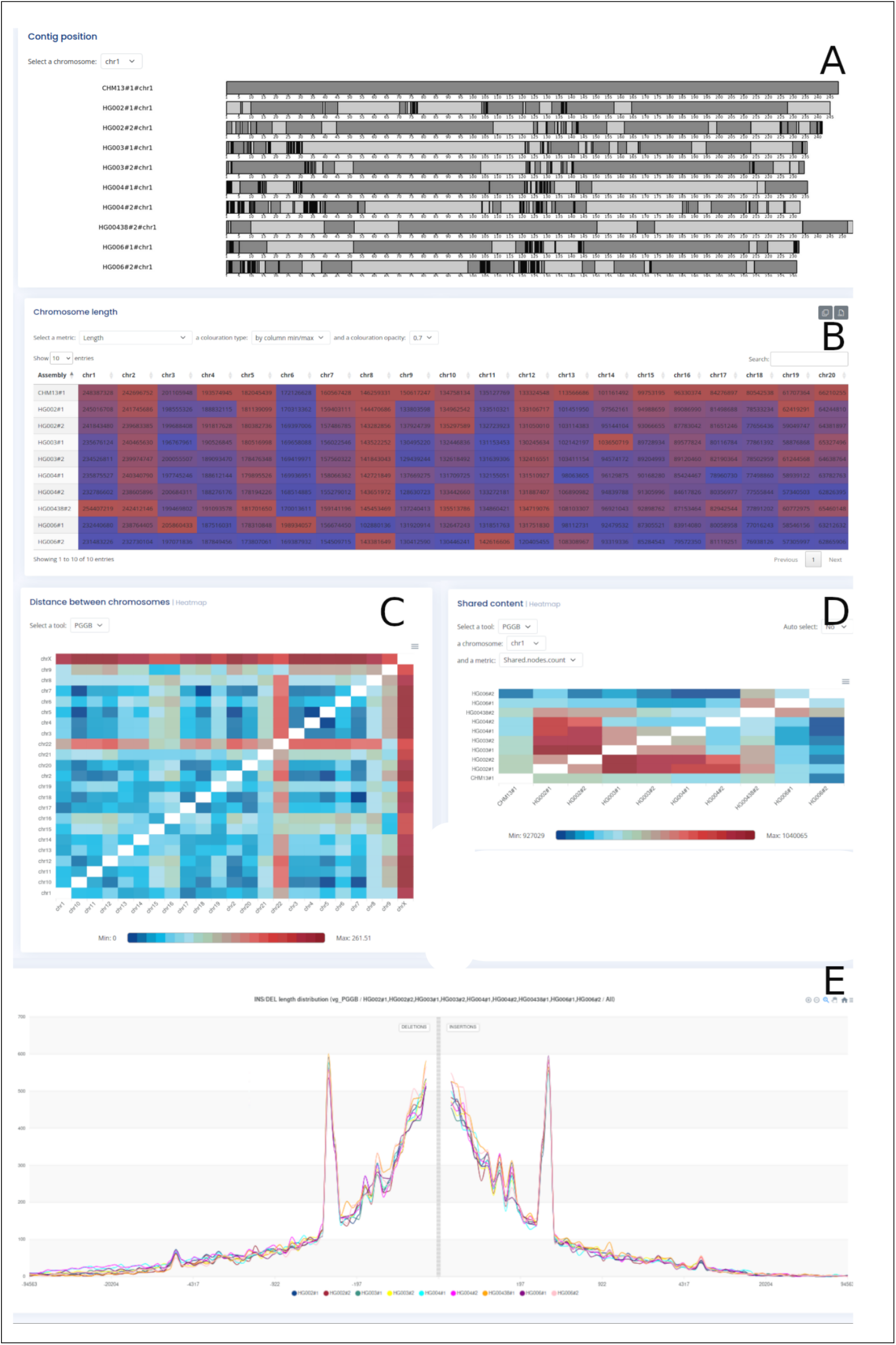
Example of `Pan1C-View` plots generated from the human dataset. (A) Contig position: displays how the contigs are distributed in assembled chromosomes. Each grey level change corresponds to a contig change. Many small joint contigs correspond to a black segment. (B) Chromosome length: heatmap showing the length of chromosomes for each assembly, can also show deviation from the mean or reference length. Chromosomes smaller than the mean length are shown within blue. (C) Distance between chromosomes: heatmap representing the euclidian distance between chromosomes pairwise based on shared content. Blue colors represents chromosomes harboring the same nodes while red color represents dissimilar chromosomes. White stands for identity.(D) Shared content: heatmap showing shared content between selected chromosomes of different assemblies. Available metrics include: shared node count, shared length, shared repeated length, shared proportion and shared repeated proportion. Tiles of the heatmap representing pairs of chromosomes sharing few nodes are within blue. White tiles correspond to self comparison. (E) Plot shows insertion/deletion variant distribution extracted from a `PGGB` graph. The horizontal axis shows the variant lengths while vertical axis represents the number of variants observed for a given length.

### 3.3. Pan1C applications

`Pan1C` was tested on three contrasted case studies, including *Homo sapiens, Triticum aestivum* and *Aspergillus fumigatus*, and representing very different types of genome complexity, ploidy level, chromosome size and various numbers of genotypes. For each dataset, we successfully ran the workflow using both `PGGB` and `MC`. Runtime, memory usage as well as other graph metrics are reported in Table 2. The wheat dataset required the largest memory resource allocation showing that genome size is a main parameter influencing computation time.

**Table 2.**
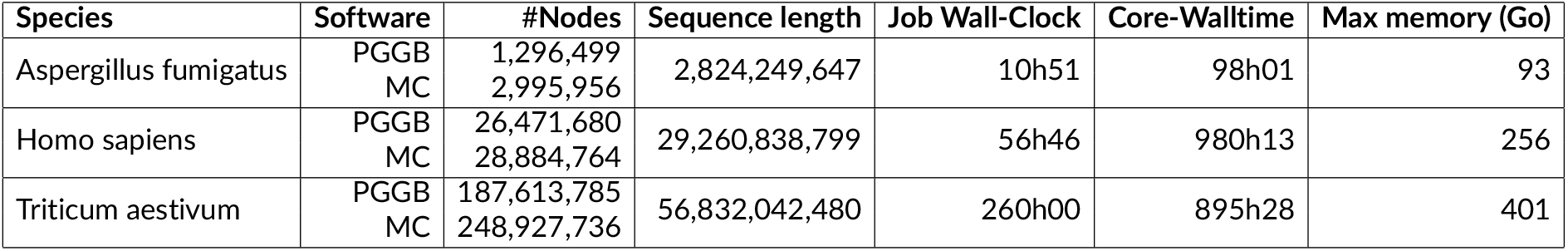
Metrics of the `Pan1C` workflow on various tests sets. ‘#Nodes’ is the total number of nodes in the final graph; ‘Total sequence’ is the sum of the per chromosome total sequence length; ‘Job Wall-Clock’ indicates the total runtime; ‘Core-Walltime’ represents the cumulative CPU time used across all cores; ‘Max memory (Go)’ shows the peak RAM usage.

`Pan1C` main output is the `Pan1C-View` archive. The reader can access the result pages using the following links for *Homosapiens, Triticumaestivum* and *Aspergillusfumigatus*.

`Pan1C-View` provides users with tables and plots that facilitate metric interpretation and comparison of `MC` and `PGGB` generated pangenomes. Some graphics are shown in Fig. 2 for the human assemblies and pangenome. Assemblies for chromosome 1 of all haplotypes and genomes are illustrated in Fig. 2-A. The CHM13 reference genome and the synthetic HG00438 (GRCh38) appear as longer sequences. This was expected as CHM13 represents the first telomere-totelomere human genome sequence, while HG00438 was constructed from multiple individuals. Other assemblies appear as shorter, fragmented, with both haplotypes of the same chromosome showing more similar sequence lengths than sequences from different chromosomes (Fig. 2-B). Heatmap views highlight how dissimilar chromosomes X and 22 are (Fig. 2-C) when compared to others. Figure 2-D clearly illustrates the extent of sequence shared between assemblies. Interestingly, both chromosomes are acrocentric, and their assemblies differ substantially in length, likely reflecting the abundance of repetitive regions. Finally the length distribution of deletions (left) and insertions (right) variants is represented in Fig. 2-E where we can clearly observe a peak at 300 bp, corresponding to the large content of Alu short interspersed elements (SINEs.)

## 4. Discussion and conclusion

We developed `Pan1C`, a unified workflow that integrates two complementary approaches, `PGGB` and `MC`, for a chromosome-by-chromosome pangenome construction. This approach, however, miss inter-chromosomal translocation events that remain poorly characterised by both `PGGB` and `MC`. Although many existing pipelines support whole-genome graph building, chromosome-level approaches are widely used in practice because they offer improved scalability while preserving biologically intra-chromosomal variations. `Pan1C` was designed to implement such a strategy and was successfully applied to three datasets representative of contrasted genome complexity and assembly quality. Its application to the large and highly repetitive wheat genome demonstrated the feasibility and relevance of this strategy for species in which whole-genome alignments are computationally demanding.

The quality of input assemblies is critical when constructing a pangenome, because tools rely directly on the assemblies to represent variation accurately. When contigs are fragmented or inconsistently labelled across genomes, `Pan1C` provides utilities within a pre-processing step to assist users in organising contigs into chromosome-scale sequences.

As a direct consequence of its design, `Pan1C` may propagate mis-located and mis-oriented contigs when scaffolding with `RagTag`. This limitation reflects a deliberate choice, prioritising computational efficiency, robustness, and simplified downstream analyses. This pre-processing step was effective for scaffolding contig level *Aspergillus fumigatus* assemblies and enabled consistent chromosome-level analyses.

By standardising preprocessing, graph generation, and visualisation steps, `Pan1C` makes direct comparisons between methods easier on identical inputs, allows for better reproducibility and aligns with emerging community best practices for pangenome analysis.

The workflow integrates state-of-the-art tools, is fully documented, includes examples and requires minimal configuration. Its user-friendly interface gives access to summary statistics, intermediate files, final chromosome-level graphs and visualisation of intra-chromosomal variations.

In this way, `Pan1C` supports researchers investigating intra-species diversity by enabling efficient, scalable and comprehensive pangenome graph construction.

Future directions include extending `Pan1C` with additional options for downstream analyses, such as `PanGenie` [7] for genotyping, `VG giraffe` [21] for read alignment, and other tools for pangenome-based annotation transfer. These extensions will further enhance the versatility of `Pan1C` as a unified framework for high-resolution, population-scale genome analyses.

## 5. Funding

This work was funded by the Agence Nationale de recherche, programme France2030’s project BReiF (ANR-22-PEAE-0014).

## Notes

### Competing Interest Statement

The authors have declared no competing interest.

### Summary of Updates

We have added a paragraph presenting other pipelines generating pangenome graphs.

